# SpatialFormer: Universal Spatial Representation Learning from Subcellular Molecular to Multicellular Landscapes

**DOI:** 10.1101/2025.01.18.633701

**Authors:** Jun Wang, Yuanhua Huang, Ole Winther

## Abstract

Spatial transcriptomics quantifies gene expression within its spatial context, significantly advancing biomedical research. Understanding gene spatial expression and the organization of multicellular systems is vital for disease diagnosis and studying biological processes. However, existing models often struggle to integrate gene expression data with cellular spatial information effectively. In this study, we introduce SpatialFormer, a hybrid framework combining convolutional networks and transformers to learn single-cell multimodal and multi-scale information in the niche context, including expression data and subcellular gene spatial distribution. Pre-trained on 300 million cell pairs from 12 million spatially resolved single cells across 62 Xenium slides, SpatialFormer merges gene spatial expression profiles with cell niche information via the pair-wise training strategy. Our findings demonstrate that SpatialFormer distills biological signals across various tasks, including single-cell batch correction, cell-type annotation, co-localization detection, and identifying gene pairs that are critical for the immune cell-cell interactions involved in the regulation of lung fibrosis. These advancements enhance our understanding of cellular dynamics and offer new pathways for applications in biomedical research.

## 1 Introduction

Recent advancements in single-cell technologies that integrate spatial information have, providing unprecedented detail, significantly improved our understanding of human diseases and biological mechanisms [1, 2]. Compared to traditional bulk and dissociated single-cell RNA sequencing approaches, image-based spatially resolved single-cell methods, like Xenium [3] and CosMx SMI [3], approach sub-cellular resolution that shed light on cell-to-cell communication [4] and the discovery of cellular niches [5]. Specifically, instead of using only gene expression, the coordinates of the transcripts and cells from these technologies make it possible to map the biological features back to the fixed tissue context and create an opportunity to capture the relative relationship of gene-gene and cell-cell pairs, and even their global correlation. Therefore, it is critical to develop methods to harness such information and resolve different biological tasks at a comprehensive scale.

Exploring cell-cell interactions (CCIs) and cell-cell communication (CCC) among different cell types through expression profiling facilitates the study of how intercellular communication contributes to cell and organ functionality. For instance, tools such as CellChat [6] and CellphoneDB [7] rely solely on expression permutation to identify significant ligand-receptor interactions. Mapping cells to their spatial context using transcriptomics and measuring the gene co-expression can physically elucidate the relationships between cells across various organs and tissues. For example, SpatialDM utilizes bivariate Moran’s statistics to detect spatially co-expressed ligand and receptor pairs [8]. However, CCIs and CCC within the tissue microenvironment are systematically regulated by multivariate factors, including cellular and subcellular effects. Specifically, CCIs may recruit different modes of CCC, leading to perturbations in gene expression levels and potentially affecting gene spatial localization, thereby altering gene-gene interaction patterns. Related tools that globally consider these factors are still unexplored. Consequently, modeling subcellular gene-gene interactions alongside cell-cell interactions can enhance the understanding of global variations in CCC and CCIs across different tissues and organs. This approach is particularly valuable in tracking disease progression and tissue development.

Gene expression profiles encapsulate intricate gene-gene co-expression and interaction insights. Unfortunately, spatial information often dissipates in conventional transcriptomic data when using single-cell transcriptomics data. Gene expression tends to be dynamic between different biological processes and environmental stimuli. For example, human neurons demonstrate autonomy by extending long processes from the cell body. This “decentralization” involves mRNA’s selective localization and translation, responding to external stimuli [9]. Furthermore, mRNA localization can facilitate protein synthesis, which is crucial to maintain or establish cellular polarity [9].

Recent advances in self-supervised learning, particularly with transformer architectures [10], have utilized extensive data sources to enhance semantic understanding in various natural language processing applications [11]. In the realm of single-cell biology, the cells can be represented by genes with different expression levels, analogue to the sentence with different words. Pre-training transformer models with large-scale single cell and single nucleus data from diverse tissues and organs, as available from HCA [12] and CELLXGENE (https://cellxgene.cziscience.com/), offering a comprehensive representation capable of querying biologically meaningful patterns and providing insights into tissue biology. Notable models include Geneformer [13], scGPT [14], Nicheformer [15], scFoundation [16], and SCimilarity [17]. However, none of these models are designed to incorporate single-cell spatial information in their pre-training schemes, as they rely solely on large-scale single-cell expression count profiles. Therefore, existing single-cell foundation models miss the key spatial contexts, lacking the power to detect cellular cooperations. Even Nicheformer, which considers both single-cell transcriptomics and spatial data, uses only the expression count matrix, omitting spatial coordination and cell-cell co-localization information. To address these limitations, we introduce a novel transformer-based model, SpatialFormer. This model is effectively pre-trained in a multitask language model scheme, leveraging spatial information by combining gene-gene and cell-cell proximity data integrated with gene expression profiles. Specifically, these multitasks encompass gene co-localization and cellular proximity information, effectively facilitating the learning of a spatially aware representation. We demonstrate that incorporating multi-scale features from multi-modalities enables the generation of richer representations. SpatialFormer is evaluated at both cell and gene levels, proving its state-of-the-art performance in cell type annotation and batch integration. Furthermore, SpatialFormer is extensively applicable for investigating cell-cell co-localization, unraveling crucial cell-cell communication signals, such as macrophage and T cell interactions in pulmonary fibrosis tissue.

## 2 Results

### 2.1 SpatialFormer model architechture

SpatialFormer is based on the SqueezeFormer architecture [18], which is recognized for its scalability and effectiveness in speech recognition tasks. By employing a transformer attention mechanism, SpatialFormer captures long-range relationships between genes and integrates this approach with convolutional layers to extract regional features (**Figure 1b**). The underlying hypothesis is that the spatial expression of genes is intricately linked to intercellular communication and intracellular gene interactions (**Figure 1c**). The SqueezeFormer architecture facilitates the identification of key aspects regarding 1) gene-gene connections at subcellular resolution and 2) spatial variations in gene-gene interactions driven by cell-cell connections across different cellular niches. This spatial awareness enhances our understanding of gene expression within various niche contexts, potentially shedding light on gene regulation under cellular perturbations, including external conditions such as disease or normal states, stimuli such as temperature, and variations related to developmental stages. In contrast to the natural language process, where the words can be encoded sequentially, the single-cell spatial transcriptomics data are inherently structured and contextually interchangeable. Therefore, novel strategies are imperative to effectively address the two-dimensional and even three-dimensional characteristics of spatial gene expression. Analogous to Geneformer [13], we utilize the ranked gene expression as our input. To incorporate multi-dimensional structural information, an ablation study was conducted to identify the most effective strategy within the pre-training framework (**Supplementary Information**). We selected the multitask learning scheme, which includes: 1) Masked Token Prediction (MT); 2) Gene-gene co-occurrences (CO) prediction; and 3) Paired-wise cell prediction (PC) (**See Methods**) (**Figure 1bd**). The CO and PC tasks are augmented by integrating gene embeddings within the embedding layers, employing GraphSAGE to pool global transcripts [19] (**Supplementary Figure 1a**), thereby facilitating accurate learning of gene cooccurrence. The PC task aims to capture the cellular co-localization features within tissue by learning from positive and negative cell proximity (**see Methods, Figure 1d**).

**Figure 1.**
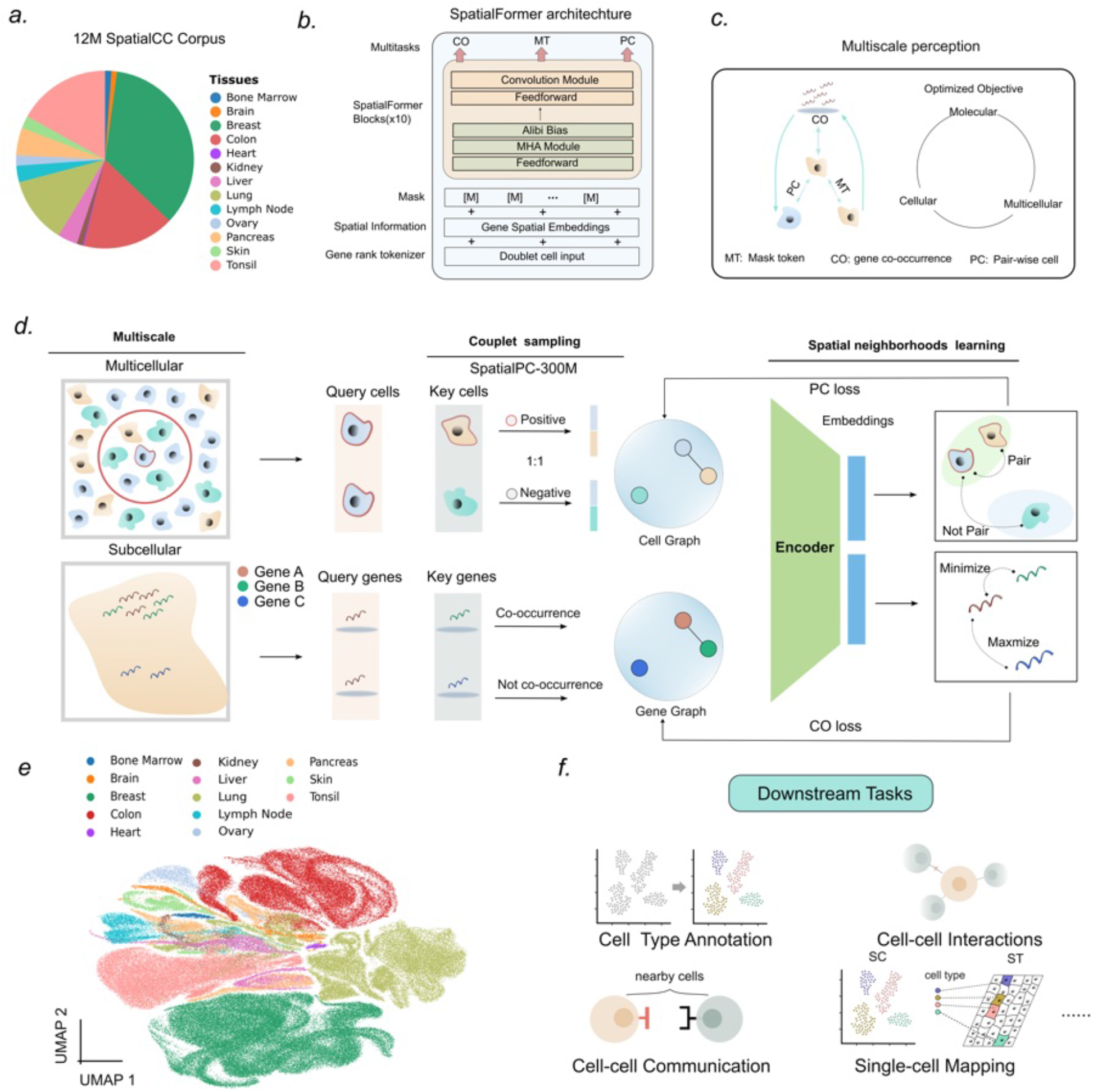
Overview of the Spatialformer architechture. **a**, The proportion of tissue types included in our 12M SpatialCC Corpus. **b**, The schematic of the SpatialFormer illustrates its structure and functionality. The ranked expression sequence is input into the SqueezeFormer unit to capture both global correlations and regional detail. The MT head is utilized to predict masked tokens, while the CO head is designed to enhance the model’s subcellular spatial awareness. Additionally, the PC head enables the model to understand whether two cells are connected within the tissue slides. **c**, Conceptual plot of the niche interaction of different scales of cell. Cell interactions can modulate gene pathways and influence gene-gene spatial localization, thereby affecting fundamental processes within the microenvironment. **d**, In the pretraining stage, two supervised learning tasks are utilized. The multicellular concept is developed by evenly constructing positive and negative cell pairs as SpatialPC-300M dataset. Molecular understanding is enhanced by building a gene-gene co-occurrence graph based on transcript communities. Both supervised tasks are integrated with the unsupervised task MT, enabling the training of the model in a multi-task learning scheme.**e**, The unlabeled pre-trained SpatialFormer embeddings are visualized using UMAP, colored by tissue types. The UMAP plot displays a random 2% subsample of the entire SpatialFormer embeddings (n = 177,452). **f**, The downstream tasks that SpatialFormer can resolved. CO, co-occurrence; MT, masked token; PC, pair-wise cells.

To ensure a diverse and comprehensive dataset for pre-training, we exclusively curated data from Xenium, which has demonstrated superior sensitivity, a wider dynamic range, and improved alignment with single-cell reference profiles compared to other popular commercial platforms [3]. Additionally, to incorporate variability across different conditions, we included healthy and disease samples from 13 distinct tissues (**Figure 1a**). This condition information can be integrated into the input sequences, instructing the model to learn the relavant conditional representations.

Following pre-training, we visualized the cell embeddings using the Uniform Manifold Approximation and Projection (UMAP) method on a 2% subsample of the pre-training data (**Figure 1e**). The UMAP visualization demonstrates that our model effectively distinguishes the intricate differences among various tissues, even without prior cell type information in the unlabeled Xenium data, by utilizing the pretrained cell embeddings. This underscores the model’s exceptional ability to capture biological variations. Afterwards, the pretrained model can be used for a variety of downstream tasks (**Figure 1f**).

### 2.2 Transcripts’ spatial co-occurrence between genes

Traditional cell markers focus solely on expression levels, overlooking spatial features and interaction dynamics. The subcellular distribution of mRNA transcripts influences their binding partners and translation rates, thereby affecting protein concentration and localization [20, 21]. Therefore, integrating gene co-occurrence with expression data is essential for accurately identifying and classifying cell types based on their spatial and functional characteristics [22]. This approach is believed to reveal additional features not captured by gene expression alone.

We modeled gene-gene physical neighbors by constructing an adjacency matrix for genes and their proximal neighbors (**See Methods**). Rather than applying global statistical filtering to co-localized genes [22], we concentrate on genes co-localized within specific regional communities (**See Methods**). This method preserves the diversity of gene distribution, which is particularly advantageous for understanding gene co-occurrence during early perturbation phases and within distinct organelles. We developed the gene co-occurrence (GCO) matrix based on two primary criteria: 1) Co-expressed genes typically share similar expression levels; and 2) Gene co-localization can be indicative even if they are not part of recognized marker lists. Notably, gene pairs with relatively low expression levels can still be classified as co-occurring and offer valuable insights for cell type identification if they are distinguishable (**Supplementary Figure 1b**).

To calculate spatial co-occurrence genes, we selected an annotated Xenium slide of human pulmonary fibrosis featuring 38 cell types (**Supplementary Figure 2a,b**) [23]. Although the differentiating ciliated markers, such as EGFR, EPCAM, and DUOXI, indicate cell type locations within the tissue, they exhibit low precision and specificity in delineating cell types (**Supplementary Figure 2c-e**). In contrast, differentiating ciliated cells can be clearly defined using gene co-occurrence (GCO) pairs as cell markers (**Supplementary Figure 3**). While individual genes within GCO pairs may not accurately represent cell types and can exhibit broad expression across the slide (e.g., XBP1 and HLA-DRA), combining them with other spatially proximate genes that also show high expression trends significantly enhances specificity and accuracy for identifying differential ciliated cell types in spatial contexts (**Supplementary Figure 2f, Supplementary Figure 4**). This observation demonstrates that employing multi-modal information—integrating gene expression with spatial proximity—facilitates more effective identification of cell types compared to traditional single-candidate lists commonly used in single-cell analyses. Furthermore, integrating gene-spatial information into self-supervised learning can substantially enhance the model’s ability to learn, providing a robust complementary approach for improving the acquisition of sequence embeddings.

### 2.3 Cell identities, niche classification and batch integration

SpatialFormer is the first single-cell language model trained exclusively on single-cell spatial transcriptomic data with small gene panels. In contrast, existing models such as scGPT [14] and scFoundation [16] have been pre-trained on extensive single-cell RNA-seq datasets featuring tens of thousands of assessed genes. To evaluate whether SpatialFormer could effectively capture cell type signals while utilizing a limited design panel of hundreds of genes, it underwent further pre-training focused solely on the masked token prediction (MT) and gene co-occurrence (CO) tasks to enhance the spatial context of gene expression. Consequently, the PC task was omitted to adapt the model for individual cell input.

We measured performance across two supervised tasks: cell type prediction and cell niche prediction, both aimed at assessing cell identities within spatial contexts (**Figure 2a-d**). A simple probe network was employed to fine-tune the model, adapting it for predicting different labels. An independent slide of pulmonary fibrosis, measured by Xenium and comprising 343 selected genes, was chosen for evaluation [23].

**Figure 2.**
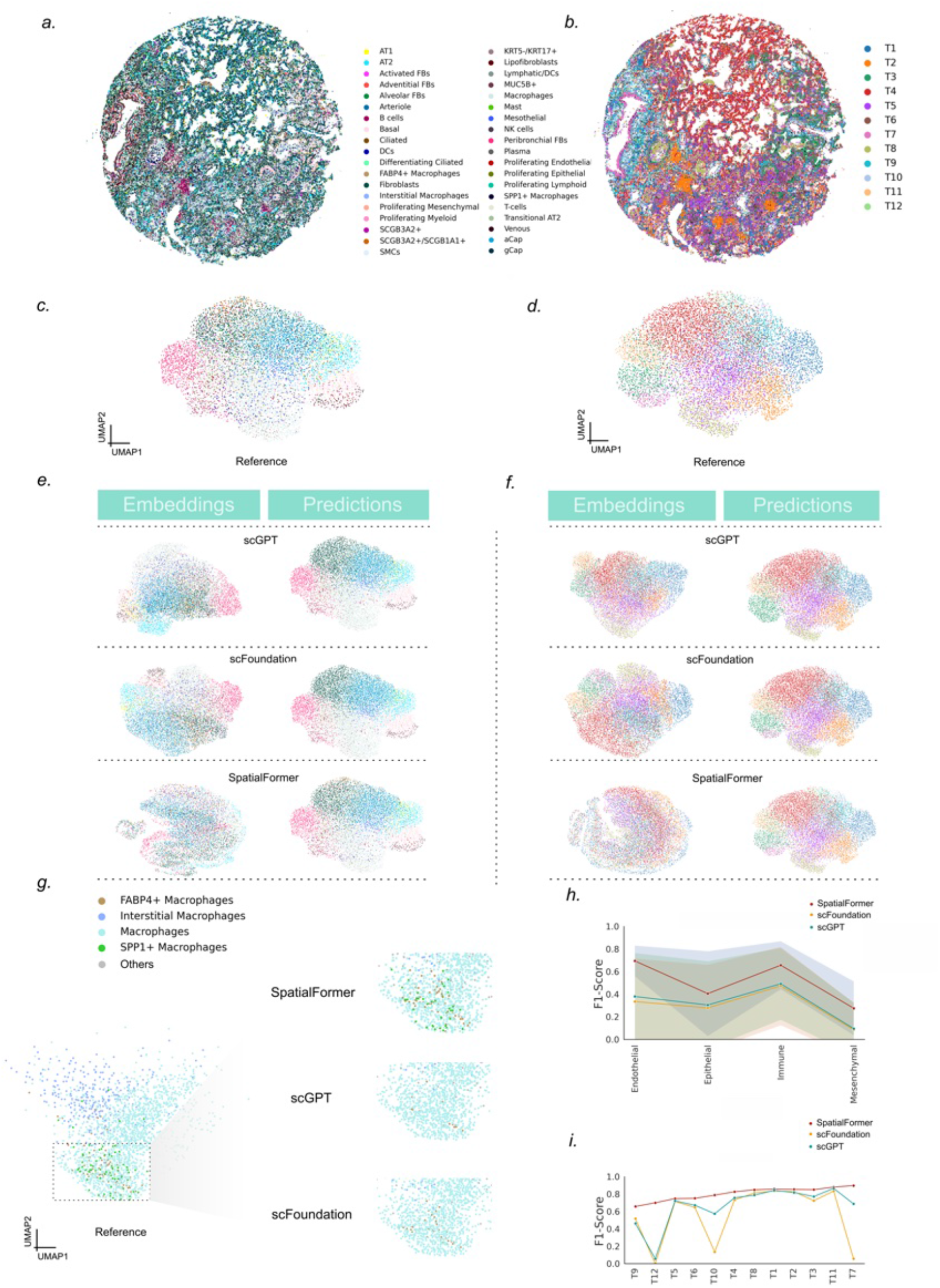
Prediction results of cell annotations. **a-b**, A pulmonary fibrosis tissue sample, named VUILD110, is depicted with cell types and niches projected onto the H&E images. **c-d**, UMAP visualizations of cell type embeddings and niche embeddings, showing 10% of the samples selected for visualization based on cell types and niches, respectively. **e-f**, UMAP visualizations of embeddings generated from different single-cell language models are shown in the left panel, while the predicted results from the probe network are displayed in the right panel. **g**, An extracted UAMP plot indicating main and subtypes of the macrophages; The left panel is the reference d1is8tribution and the right panel shows the prediction results of the SpatialFormer, scGPT, scFoundation, respectively. **h**, The line plot evaluates the performance of different language tools across various lineages, with shaded areas representing the range of F1 score across cell types within specific lineages. **i**, A line plot illustrating the performance of cell niches across three language tools, with colors consistent with those used in panel h.

Beyond the exclusive use of expression data, the embeddings generated by SpatialFormer display greater diversity compared to those from scGPT and scFoundation, particularly regarding cell types and niches (**Figure 2e,f**). Specifically, endothelial and immune cell types exhibited higher accuracy than the epithelial and mesenchymal lineages as 0.70 and 0.68, respectively (**Figure 2h**). SpatialFormer consistently outperformed both scGPT and scFoundation in predicting macrophage subtypes. It demonstrated a significantly better performance in identifying FABP4+ macrophages, achieving an F1 score of 0.71 compared to F1 scores of 0.31 for scGPT and 0.50 for scFoundation. Notably, the other two foundation models were unable to recall any SPP1+ macrophage cells in their predictions (**Figure 2g, Supplementary Figure 5j**, recall = 0), while SpatialFormer successfully recalled 20 SPP1+ macrophages, yielding a recall of 0.27 (**Supplementary Figure 5**). SPP1+ macrophages are rarely detected in control lung tissue, and their increased expression has been associated with exacerbated pathology. These findings suggest that SpatialFormer has potential utility in predicting rare cell types that play a significant role in disease progression. In terms of niche performance, SpatialFormer consistently outperformed others, peaking at T7, which corresponds to the transitional epithelial (**Figure 2i**). These results suggest that the richer representation of SpatialFormer is more effective at learning the detailed features of subtypes, which other single-cell independent models fail to capture.

The predictions of cell types validate the e”cacy of our model in utilizing a limited list of genes to discern cell identities within the spatial dataset. Additionally, we applied the model to two single-cell datasets, including batches collected from COVID-19 (18 batches) [24] and the kim lung dataset (14 batches) [25]. Our model outperformed scGPT in batch correction in the kim lung dataset and showed comparable performance in the COVID-19 dataset (**Supplementary Figure 6**,**7**).

These results collectively indicate that SpatialFormer, even when pre-trained on a small gene panel, can capture critical gene signals to make accurate cell type predictions. Furthermore, these signals are su”cient for distinguishing different cell types and capturing invariant batch information, demonstrating the model’s generalization capabilities and robust predictive power when utilizing multimodal information for single-cell analysis, including gene expression and spatial context.

### 2.4 Cell-cell co-localization analysis

Tissue function is sustained through the coordinated activities of various cell types [26]. The patterns of cell-cell co-localization can reveal insights into tissue organization under different developmental and disease conditions. For instance, in cancer, increased cell density may occur due to the recruitment of immune cells within the tumor microenvironment [27]. This dynamic composition can lead to changes in interactions, where the query cell engages with different groups of cell types in response to environmental shifts. Therefore, accurately predicting the co-localization of query cells and their neighbors holds significant prognostic value.

SpatialFormer is pre-trained using a pair-wise approach, enabling the model to effectively capture critical pair-wise features associated with cellular niches within the microenvironment by contrasting positive and negative pairs. By fine-tuning the model on slides exhibiting representative cell type colocalization patterns, we can seamlessly transfer the learned cell-cell interaction features to a new tissue context, allowing for direct inference of cell-cell co-localization solely from expression profiles in single-cell data (**Figure 3a**).

**Figure 3.**
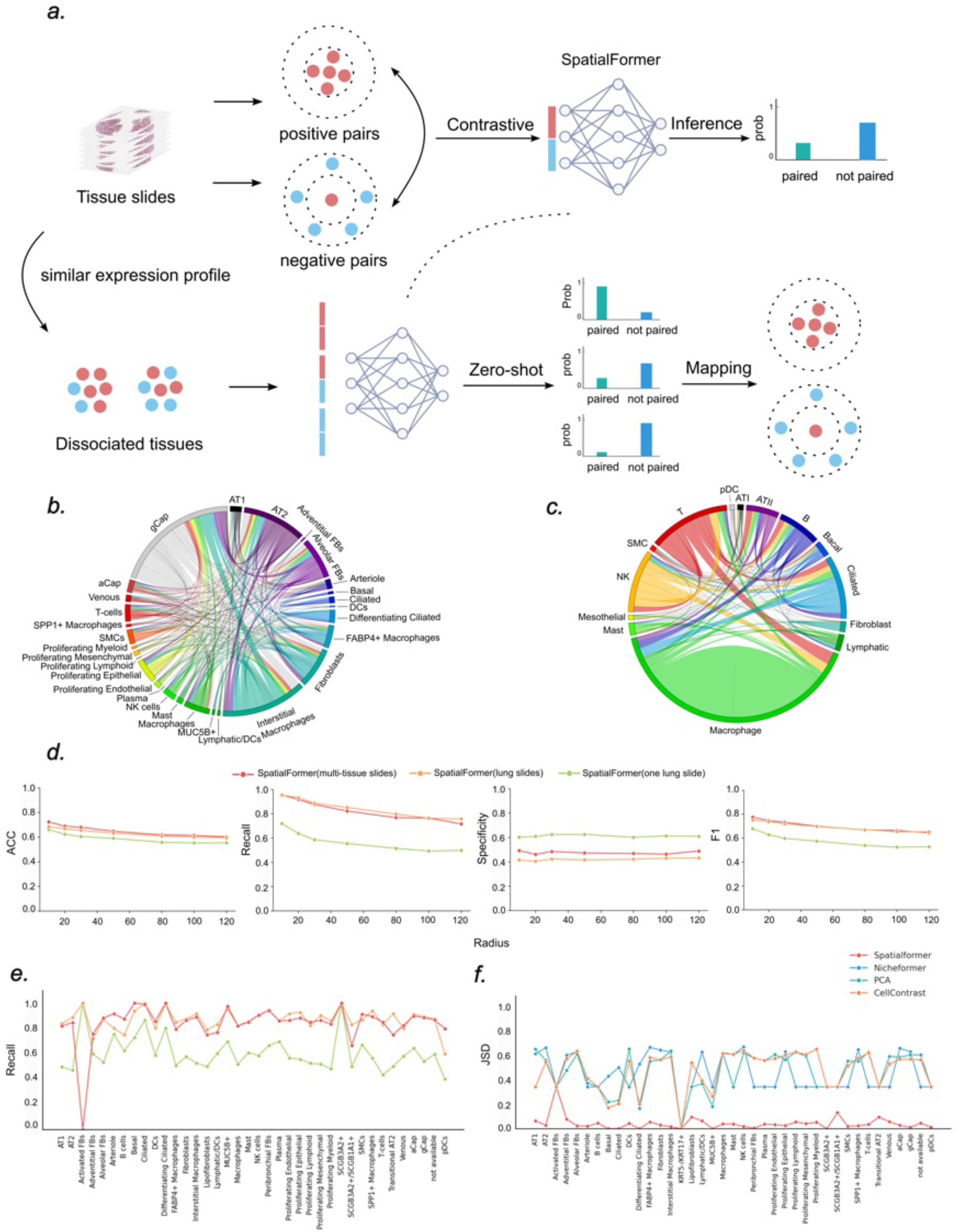
SpatialFormer Co-localization performance. **a**, The schematic plot illustrates the prediction of cell-cell co-localization. SpatialFormer is trained on tissue slides using contrastive learning to differentiate between proximal and distal cells. Subsequently, the fine-tuned model can be applied to single-cell data lacking spatial information for predicting cell-cell co-localization. The accompanying Hematoxylin and Eosin (H&E) image depicts pulmonary fibrosis, specifically from slide THD0008. **b**, visualization of cell-cell co-localization in lung slide THD0008 among different cell types, with connected cells comprising fewer than 500 excluded for clarity. **c**, The circus plot displays the predicted co-localization of cell types, highlighting the selected cell types that overlap with the spatial annotations.**d**, performance metrics for cross-slide prediction in identifying cell neighbors, including accuracy, recall, specificity, and F1 score. **e**, visualization of t1h9e recall scores for all cell types in slide THD0008. **f**, The Jensen-Shannon divergence illustrates the distribution of all cell types across slide THD0008, bench-marked against three other generated embeddings. Lower values indicate better performance.

In this study, we utilize SpatialFormer to transfer co-localization information related to lung fibrosis from Xenium spatial data to single-cell RNA-seq data from Taylor et al. [28]. We only select those cell types that overlap between the single-cell and Xenium datasets. The zero-shot results indicate a strong co-localization of immune cell types, which is similarly observed in the SpatialFormer predicted results based on the expression profiles (**Figures 3bc**). This finding suggests that SpatialFormer can potentially be applied in scenarios where paired spatial data are unavailable by extrapolating information from expression profiles.

Unlike brain tissue, where cellular composition is conserved and consistently connected to their functional neighbors, the cell-cell composition in unstructured tissue slides, such as lung tissue, can vary significantly. This variability complicates the ability to learn a globally generalized cell-cell co-localization feature across all slides. To assess our model’s effectiveness in capturing common features of cell-cell interactions within tissue, we pretrained SpatialFormer across three different settings: 1) multi-tissue slides; 2) one similar slide (VUILD102LF); and 3) Series of similar slides. We conducted evaluations using a leave-out lung test disease slide (THD0008) and found that the pre-trained model on all tissues (61 slides) achieves the best performance. Namely, the more general spatial expression feature we learned, the better performance we can get in the test sample. Conversely, only the fundamental features influencing cell-cell interactions can be generalized across different slides under similar conditions. Notably, recall peaked at 0.95 when using the multi-tissue pre-trained model to do directly inference at the 10 *µm* resolution (**Figure 3d**). Furthermore, the large-scale pre-trained model slightly outperforms the lung tissue-specific model at a tissue scale for capturing cell-cell co-localization information, achieving a maximum F1 score of 0.77, representing a 15% improvement over the single-slide model (F1=0.67) and 3% improvement over the tissue scale model (F1=0.75)(**Figure 3d**). Therefore, pre-training on a large scale of cell-cell co-localization data may have advanced the model’s ability to learn global generalized features with the diversity of cell types under variant spatial contexts. All models struggled to accurately predict cells that were physically distant from the query cells, indicating that the spatial features learned from the expression profile may diminish when cells are far apart (**Figure 3d**). Despite these challenges, SpatialFormer successfully predicts cell types with significant spatial distribution in 30 *µm* resolution even in the nonstructural lung samples (**Figure 3e**). For instance, ciliated cells, which exhibit pronounced spatial patterns on the slide (**Supplementary Figure 2A**), were predicted with a recall near 1 for multi-tissue model and got a recall of 0.9 even training on just one related slide.

Furthermore, we compared SpatialFormer with three other methods for identifying cell neighbors on the same slide of pulmonary fabrosis. Unlike SpatialFormer, NicheFormer [15] focuses solely on spatial expression profiles. Additionally, we included CellContrast [29], a dedicated model known for its state-of-the-art performance in identifying spatially proximal cells through contrastive learning. We evaluated these methods by measuring the closest cell proximity using cosine similarity across all candidate cells on the slide. Our findings reveal that, despite claims of encoding spatial information, NicheFormer failed to accurately capture cell-cell co-localization patterns for each cell type, resulting in random proximity predictions. CellContrast also demonstrated limited predictive abilities in lung tissue, even though it shows strong performance in structured tissues like the brain (**Figure 3f**).

### 2.5 Perturbation identifies interaction-associated gene pairs

The cellular niche operates as an ecosystem profoundly influenced by the ions, metabolites, integrins, and proteins secreted by neighboring cells. Cell-cell communication can activate downstream signaling pathways through cognate receptors, ultimately leading to changes in transcription factor activity, which in turn affects gene expression levels. Therefore, the dynamics of cell-cell interactions can be understood through gene expression levels [26]. Notable examples include CellTalker [30], CellChat [6], and Cell-PhoneDB [7]. However, there is currently no communication score computed from the covariant factors that takes into account the integrated factors in the cellular niches, including all genes rather than only ligands and receptors. Here, we aim to use SpatialFormer to investigate all the mediators for cell-cell communication within the niches.

In the progression of pulmonary fibrosis, granuloma formation is a critical characteristic for both diagnosis and treatment. Macrophages and T cells are found co-localized in this region with unprecedented strength (**Figure 4a**). However, the genes that regulate these cell-cell interactions remain elusive. We employed SpatialFormer to conduct Iterative Gene Pair Perturbation (IGP) (**See Methods**) to identify significant gene-gene pairs in the granuloma region by screening a vast number of possible perturbations at the scale that would be impossible in the wet lab. Specifically, genes are iteratively knocked out in the pair-wise cell paradigm, with those genes exhibiting significant effects on the predicted logits considered important for regulating cell-cell interactions. As a result, 605 pairs were found to significantly contribute to cell pair-wise interactions (**Figure 4b, Supplementary Table3**). Importantly, these gene pairs do not exhibit a significant correlation with gene rank, indicating that the signaling of these pairs is not dominated by highly expressed genes (**Figure 4b**); lower-expression genes can also play a vital role in regulating these interactions. Notably, the significant pairs are associated with the regulation of collagen-containing extracellular matrix (ECM), which is crucial for lung structure and function, aligning with the pathogenesis of fibrotic diseases (**Figure 4c**). Furthermore, many of these paired genes are linked to immune system functions, further confirming the concentration of immune cells within the granuloma (**Figure 4d**).

**Figure 4.**
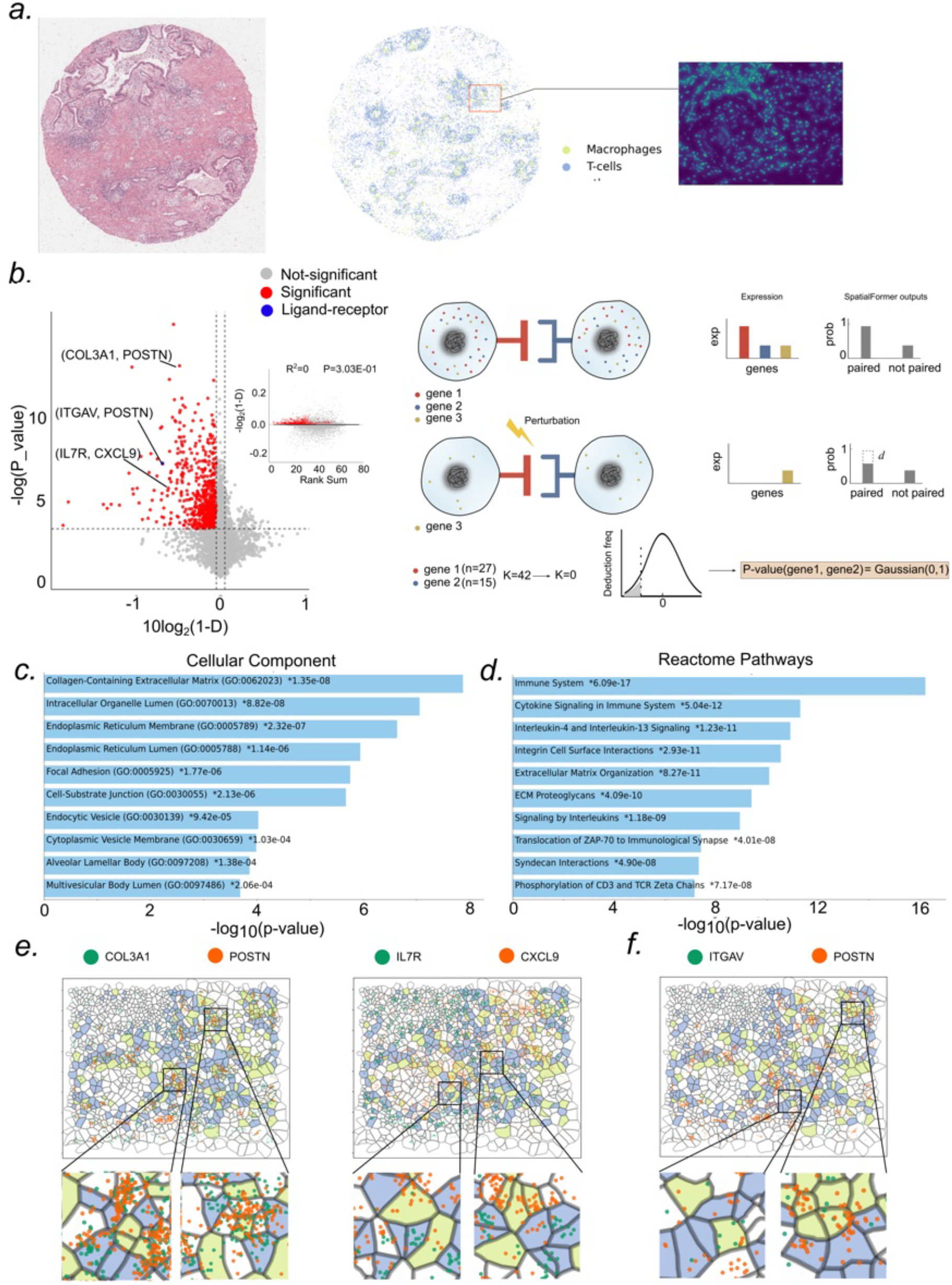
Gene-Gene perturbation analysis reveals key cell-cell interaction features. **a**, the left image displays a Hematoxylin and Eosin (H&E) stain of pulmonary fibrosis from slide VUILD102LF. The middle plot illustrates the distribution of macrophages and T cells across the slide, represented in distinguishable colors. The square subplot on the right provides a zoomed-in view of the pulmonary 20 fibrosis morphology, highlighting the Granuloma feature. **b**, The volcano plot displays the selected significant gene pairs in red. The right panel illustrates a cartoon representation of the in-silico perturbation. Paired genes are depleted, and the importance of each gene pair is quantified by the predicted logits. A one-tailed t-test was conducted to assess the significance of these pairs in reducing the predicted logit mean value. The variable *d* represents the deduction value of the predicted logits. **c-d**, enrichment analysis of the top influential gene pairs in terms of gene ontology and cellular components. Panels **e** and

Finally, we assessed whether the perturbation analysis could identify canonical ligand-receptor pairs within our gene list. One ligand-receptor pairs were found to overlap significant gene pairs in the granuloma region. Significant pairs, such as *ITGV A− POSTN*, demonstrate relative light co-expression in macrophage and T-cell niches according to their low expression level but can still be predicted by perturbation analysis, indicating their importance in cell-cell colocalization of granuloma region (**Figure 4f**). This gene pair contributes to the development and progression of lung fibrosis by cell adhesion, migration, and tissue remodeling. In addition to known gene pairs, *COL*3*A*1 −*POSTN* and *IL*7*R −CXCL*9 are identified as novel gene pairs important for cell-cell interactions (**Figure 4e**). These novel pairs are recognized for their roles in inflammation, immune response, and ECM dynamics, which are biologically significant in the pathogenesis of pulmonary fibrosis. A complete list of gene pairs can be found in **Supplementary Table 3**.

Unlike traditional single-cell and bulk RNA-seq datasets, Spatial transcriptomics technologies, e.g., Xenium, offer a remarkable advantage in capturing intricate details of cell-to-cell crosstalk by verifying the presence of communication factors in situ. This capability greatly enhances the reliability of computed CCIs, providing a more accurate depiction of cellular communication landscapes. SpatialFormer introduces a novel tool designed to probe the potential transcriptomic factors that regulate CCC within subcellular-resolution spatial transcriptomics data. Researchers can delve deeply into specific Regions of Interest (ROIs), uncovering the precise gene pathways involved in proximal cell-cell interactions. This granular level of analysis empowers researchers to better understand disease processes at a cellular level, potentially facilitating the exploration and identification of novel therapeutic targets.

## 3 Discussion

We present SpatialFormer, a large-scale pre-trained single-cell foundation model that maintains spatial awareness by harmoniously modeling cells and genes within their spatial context, as well as coordinating these two modalities. By combining self-supervised and supervised learning in a multi-task learning design, we successfully integrate spatial information into the model’s hidden representations, producing richer feature representations. The generated embeddings, encompassing genes, cells, and even pairwise cell relations, enable the model to be applied in diverse downstream applications, especially the cell-cell co-localization and communication that the other language cannot achieve.

It is noteworthy that SpatialFormer is capable of not only addressing spatial tasks through the measurement of cell-cell co-localization pairs but also learning individual cell representations. Both functionalities are essential for understanding the characteristics of single cells and the pairwise gene interactions that influence cell-cell co-localization. Utilizing perturbation analysis, we can identify significant gene pairs that are co-expressed and critical in determining cell-cell interactions (CCIs). This approach can be applied to any combination of cell types, allowing for the investigation of biologically relevant ligand-receptor gene pairs as well as novel, meaningful gene pair interactions. A focused examination of spatial details can enhance our understanding of molecular perturbations, offering potential applications in drug development for various diseases.

However, the spatial transcriptomics dataset may be limited due to the restricted number of genes in pre-designed panels. Currently, we mainly measure canonical cell markers and biologically relevant genes, which can lead to missing complex gene signals critical to biological pathways. Our support extends to approximately 2000 genes across 13 human tissue types. As more spatial transcriptomics data are released, we anticipate the realization of universal gene panels that allow for the measurement of a transcriptome-wide range of genes, akin to single-cell data, thereby enabling comprehensive assessments of cells within their spatial contexts. Furthermore, with the ongoing release of non-human spatial data, cross-species models can be developed as we reach su”cient data scale.

Despite the fact that transcript co-localization is designed to enhance gene regulation insights, the physical locations of transcripts—such as intranuclear, perinuclear, and cytoplasmic—are often overlooked. Incorporating this spatial information can improve the understanding of gene ontology from the nascent state to the post-transcriptional state. However, care must be taken when integrating this data, as transcripts exhibit significant dynamic behavior in vivo. This means that transcripts can localize to various compartments in response to distinct cellular conditions, influenced by disease states, development state, or external stimulus. Therefore, integrating a temporally aware transcript model could provide a more comprehensive representation of the effective factors influencing cell-cell interactions.

In the future, we believe that an increased volume of data will significantly enhance model performance. To mitigate the constraints associated with single-modality data, integrating diverse data types in the pre-training scheme—such as multi-omics and gene sequence data—may be advantageous. Additionally, given that spatial contexts can vary dynamically throughout the tissue lifecycle, gene-gene interactions and cell-cell interactions are expected to change over time during tissue development and disease progression. There is a pressing need for a universal multi-modality single-cell spatially resolved foundation model.

## 4 Methods

### 4.1 Data collection

To keep the diversity of the input data and allow the model to learn the intricate semantic representation, we collect the unlabeled Xenium dataset with 12,000,000 cells denoted SpatialCC-12M consisting of 62 human FFPE (formalin-fixed, para”n-embedded) samples and 13 tissue types. We believe that perturbed cells can also favor the learning of the richer semantic representations, so we incorporate cells from both healthy and disease conditions. The gene panels of 10X Genomics were chosen to accurately type cells, and identify select immune, proliferation, and tumor markers. When incorporating gene panels of all the 13 tissue types, we employ the overall number of 1922 genes as our gene vocabulary, and 1.4 × 10^9^ transcripts have been measured and used for calculating the gene co-occurrence (**see 4.3**). Most of the samples use a similar number of genes in the assayed panels, and the measured transcripts per gene are comparable in both normal and disease cells (**Supplementary Figure 8de**).

Within SpatialCC-12M, we also collect latest released labeled Xenium data from lung samples with about 820,000 query cells [23]. Details of the collected dataset are provided in **Supplementary Table 1**. The dataset comprises 25 lung tissue samples, each measuring 3-5 mm, collected from 6 unaffected donors and 13 donors with pulmonary fibrosis (PF). On average, each cell in the dataset contains 69 genes.

### 4.2 Data preprocessing

To ensure the extraction of su”cient signals and to minimize noise that could affect our findings, we filtered out all cells with fewer than 30 transcripts. Each transcript was required to have a quality score of at least 20 to ensure the robustness of the fluorescence signal. In addition, genes labeled “unassigned”, “negative”, and “blank” were excluded from the analysis. To mitigate the effects of housekeeping genes, which are not typically distinguishable for cell type identification, we normalize gene expression levels by the technical median expression level as in the Geneformer [13]. This normalization process ensures that distinguishable genes associated with specific cell types receive a higher ranking, while highly expressed housekeeping genes are assigned a lower rank **(see 4.6 for more details)**. Gene expression levels were normalized to a total count of 10,000, and cells with fewer than 10 expressed genes were filtered out.

The pre-processed data are stored within the Huggingface datasets structure [31], which is based on the Apache Arrow format that allows processing of large datasets with zero-copy reads without memory constraints. The dataset can be easily downloaded for model training and inspection.

### 4.3 Gene-gene co-localization

We hypothesize that the co-localization of transcripts is nonrandom, indicating that genes that located in close proximity may exhibit similar functional trends in cell division, differentiation, and polarization [32]. SpatialFormer leverages gene-gene co-occurrence information by meticulously curating and dissecting them from the collected datasets.

To obtain gene-gene co-occurrence information, we defined *r* as the radius of a circle centered on a transcript at position (*u, v*). *r* is set to 5 *µm* for the Xenium data to ensure the robustness of the cooccurrence signal. The co-localization domain *D*(*u, v*) is the set of transcripts constrained to be within this circle:

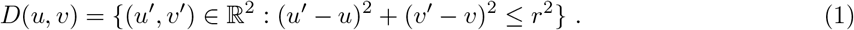

Afterwards, the connected (co-localized) transcripts were clustered using the Louvain algorithm and the transcript connection communities 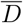 were constructed. We established a specific criterion to define genegene interactions within these communities. To ensure a strong co-localization signal, we only consider genes as co-localized if at least three copies are present within each community. For example, the cooccurrence of genes A and B is encoded as “1” if each gene exists at least 3 spots in the community 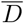.Otherwise, they should be encoded as “0”, denoting non-co-occurrence. The gene-gene co-localization information will be encoded as the matrix *G* ℝ^*m*×*m*^. After pre-processing, we make sure at least one co-occurring gene exists for each cell. Additionally, those genes with co-localization tend to have higher expression rank when compared with non-co-localized genes (**Supplementary Figure 8c**).

### 4.4 Spatial-aware gene embeddings from GraphSAGE

To enhance the gene-gene co-localization prediction, a global gene distribution embedding is generated, which is pre-trained from the subcellular distribution of the transcript neighbors.

Specifically, the global gene-gene interaction can be represented by building the computational graph without cell boundary. It is constructed using the individual transcript as the node and the edges drawn between two nodes if they are smaller than 3 *µ*m. To save memory, we randomly select 20000 cells for each slide to get the sub-graphs. We sample the samples into 5000 randomly selected “root nodes” to build the joint graph across all the samples. Then, we implemented a two-hop graph neural network using GraphSAGE to learn the spatial transcript embeddings in a self-supervised manner. The training objective is set to distinguish positive pairs and negative pairs between two transcripts. The training is stopped as the training accuracy reaches 0.82, which can be used as high-fidelity transcripts spatial embeddings. Finally, the gene spatial embeddings were calculated by averaging the spatial transcripts of the same gene across all joint graphs [23].

### 4.5 Spatial-aware cell communications

In addition to gene-gene co-occurrence at the subcellular level (as previously discussed), cells can communicate with their neighboring cells through ligand-receptor signaling, which may, in turn, influence subcellular gene distribution.

In this study, we independently compute the cellular adjacency information for each tissue sample, resulting in a distinct graph for each sample that captures the sample-specific cell-cell communications. Specifically, a centroid cell can be treated as a query cell with the connection of all the neighbors within a certain radius. To determine the optimal radius for constructing the connection graph, we assess candidate distances of 20 *µ*m, 30 *µ*m, and 40 *µ*m. Among these options, a radius of 30 *µ*m is chosen as the optimal parameter for building the cell graph, as it effectively balances Type I and Type II errors compared to the other candidates (**Supplementary Figure 9**). Afterwards, a binary sparse adjacency matrix *A* can be made to describe the overall cell-cell interaction within each tissue slide as *A* ∈ ℝ^*n*×*n*^, with *n* representing the number of cells present in the slide, *a*_*ij*_ = 1 if *d*(*x*_*i*_, *x*_*j*_) ≤ *θ*_*r*_; otherwise, *a*_*ij*_ = 0, where *d*(*·*) represents the Euclidean distance between two cells.

To enable the model to learn which features in connected cells are important within the cell niches, we randomly selected an equal number of negative pairs, where *a*_*ij*_ = 0, for each query cell *i* with *a*_*ij*_ = 1 from any key cell *j* (*j*≤ *n*). This approach ensures an equal distribution of positive and negative pairs, allowing positive pairs to occur 50% of the time.

Importantly, unlike natural language, cell pairs do not have a specific order; therefore, Both forward and backward order are included to encapsulate the interchangeability of the cell pairs. Ultimately, this results in the generation of 300 million cell pairs as the SpatialPC-300M dataset.

### 4.6 Cell expression profile represented as token sequence

The gene expression vector *x*^*i*^ for a given cell *i* can be measured on different scales using various protocols. Encoding the raw expression levels uniformly is challenging due to the varying magnitudes of these values [33]. To address this, we use relative expression levels, ensuring each input vector *x*^*i*^ has a specialized order.

Specifically, we compiled a set of genes from the collected dataset as the set of gene vocabulary with 1922 unique genes. Each gene is assigned a unique identifier based on its alphabetical order, denoted as id(*g*_*j*_). These identifiers form the vocabulary of effective tokens, serving as inputs to SpatialFormer. Thus, the input cell vector *x*^*i*^ is represented by a subset of the vocabulary set *V* ∈ ℝ^*M*^.

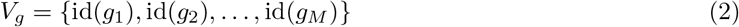

where *M* indicates the maximum length of the vocabulary.

Distinct cell types are characterized by their differential gene expression levels, reflecting their unique cellular lineages [34]. Therefore, representing cells by the order of gene expression helps encode cellular diversity. For instance, two cells of the same type may share similar expression profiles, meaning 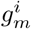 and 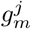 in two different cells may both have high expression levels. Similar to how word order can alter sentence meaning, we encode cell identity by ranking expressed genes in descending order. To mitigate platform-specific biases in gene measurement [33], gene expressions are normalized by their technology-specific median score. Here, we only consider the Xenium technical mean, and it can be extended for more spatially resolved transcriptomics techniques. The normalized cell vector *x*^*i*^ can be denoted as:

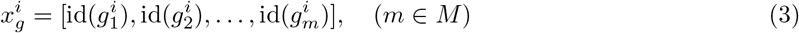

where *m* is a ranked subset of the expressed genes from the vocabulary *M*.

To further represent cell identity, we incorporate meta-information into the completed input vector *X*^*n*^. This includes biological context and protocol details, such as `<SP>`, `<TI>`, `<CO>`, and `<AS>`. The first three meta tokens encode biological context across three levels: species (e.g., “Mouse” or “Human”), tissues (e.g., “lung” or “brain”), and conditions (e.g., “disease” or “healthy”). The fourth token represents assay types (e.g., “Merfish” or “Xenium”). Incorporating these meta tokens into *x*^*i*^ enables the model to learn context-based embeddings during self-supervised learning. These four context-based tokens are placed at the beginning of *x*^*i*^.

Because none of the cells contain more than 250 expressed genes in all the collected Xenium datasets (**Supplementary Figure 8b**), we can set the maximum input length of the tokens for each cell as 500. Because we feed the pair-wise cells to make the model get the sense of their niche structure, the paired cells should have a max 500 length of input. The paired cells in the positive or negative group are separated by `<SEP>` token. Therefore, we set a fixed input length of *L* = 500 to facilitate batch processing. If a cell *i* has fewer genes than this threshold, we pad the input to *L* using the padding token `<PAD>`. The `<CLS>` token is placed as the start of the sequence. Consequently, the processed input vector *X* ∈ ℝ^*N* ×*L*^ is obtained as:

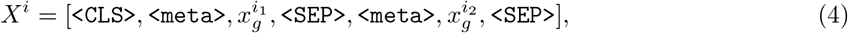

where *i*_1_ and *i*_2_ represent the first and the second paired cell, respectively.

To incorporate the gene with the global distribution context, *X* is added by the pre-trained embeddings that are trained by the GraphSAGE model (see 3.5). These embeddings can be used as the default gene spatial distribution, which can be tunable during the pre-training to fit the model in different training objectives. Additionally, *X* also adds token type embeddings to distinguish the first and the second cells in the selected cell pairs.

### 4.7 SpatialFormer architecture

SpatialFormer is a hybrid convolution-attention transformer architecture that has shown success in speech recognition tasks [18]. This model comprises *L* stacked squeezeformer blocks, each consisting of a convolutional sub-module followed by a multi-head attention sub-module. The encoded input vector *X*^*i*^ ∈ ℝ^*m*×*D*^, where *D* represents the hidden dimension of the embeddings. For a multi-head attention block:

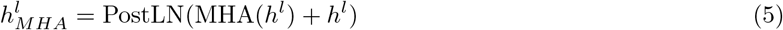

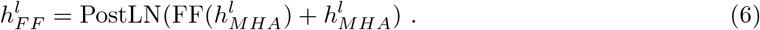

Where PostLN and FF denote the Post-Layer Normalization and feedforward layer, respectively. The MHA() denotes the multi-head attention layer, which enables the tokens to focus on the essential context during the training process. Specifically, the query (*Q*), key (*K*), and value (*V*) are linear projected by *W*_*q*,*k*,*v*_

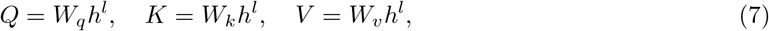

where (*Q, K, V* ∈ℝ^*m*×*D*^), and *W*_*q*_, *W*_*k*_, *W*_*v*_ ∈ ℝ^*d*×*D*^ are the trainable weight matrices for the queries, keys, and values, respectively. Where *W*_*q*,*k*,*v*_ ∈ ℝ^*d*×*D*^ denotes the trainable weights and *h*^*l*^ is the hidden representation of the *lth* specific layer. For a convolutional block:

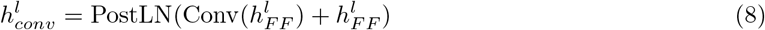

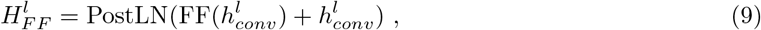

where each sub-module is connected by residues and follows the same rule of normal transformer architecture [35].

Instead of applying the positional encoding in the input token layer, we combined the squeezeformer architecture with the attention with linear bias (Alibi) [36], which is initially employed to replace the positional embeddings at the bottom of the network and extend the input sequence into longer scale. Specifically, the attention sublayer computes the attention score for a specific position *n*, setting the *n*-th query *q*_*n*_ ∈ ℝ^1×*d*^, where *d* is the head dimension. Given the set of *n*

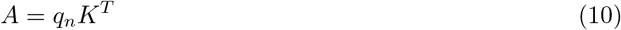

We also set the unlearnable bias P to encode the positional information.

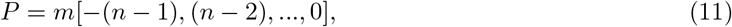

where scale m is a fixed number for different attention heads, following slope as

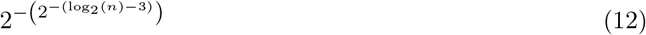

Finally, we integrate these biases together in the attention score calculation

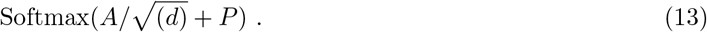

In the training and inference step, SpatialFormer doesn’t need to add the positional encoding layer and will generate the according embeddings with the gene expression profile provided.

### 4.8 Pre-training objectives

To develop a rich shared representation and enhance both learning e”ciency and accuracy, we employ a multitask learning framework for our training objectives. These objectives encompass masked token prediction (MT), spatial co-occurrence prediction (CO), and paired cell prediction (PC). The MT task is further augmented by incorporating the gene and cellular spatial context(**see 4.3 and 4.5**), enabling the model to learn low-dimension cellular representations in a spatial awareness manner by both observing gene distribution in the subcellular resolution and cell-cell interactions in the cellular micro-environment. When optimizing the MT task, the model needs to guess the masked tokens based on the revealed tokens in the sequence context. Specifically, the final output of the feedforward layer 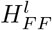 is stacked with a multilayer perceptron network (MLP) to estimate the predicted masked gene probabilities. For the specific cell *i*, the loss is calculated by the binary cross-entropy in all masked genes as ℒ_MT_:

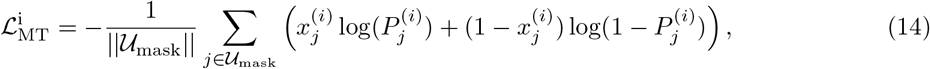

where *j* denotes the randomly masked genes, and ||𝒰_mask_|| represents the number of masked genes. The robust filtered spatial co-occurrence matrix is introduced to learn the subcellular distribution of the genes, which can be defined as a supervised classification task by optimizing the binary cross-entropy ℒ_CO_.

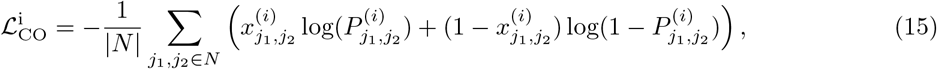

where *j*_1_, *j*_2_ represents the index of two genes, and |*N*| is the total number of spatially connected gene pairs. 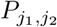 is the predicted pair probability of 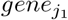 and 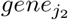.

In the PC task, the objective is easily built as to whether two input cells are a pair or not, which indicates whether they are spatially close to each other and may have potential cell-cell communication. The learned embedding of the `<CLS>` token is used to represent whether two pair cells potentially communicate with each other. The loss can also be defined by binary cross-entropy as ℒ_PC_

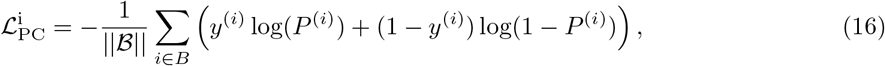

where ||ℬ|| denotes the batch size and *P* ^(*i*)^ represent the probability of the interaction between two cells. Because we build the positive and negative pairs evenly, the positive and negative cell pairs should be around 1:1 in a certain batch.

It is important to note that the mixed supervised and self-supervised classification tasks render our training objective an optimization problem in the context of multitask learning. When optimizing the multitask loss function ℒ_MT_, the model is required to learn an abstract, high-dimensional representation to infer masked gene tokens based on the contextual information provided by other genes, which are ranked in descending order. The learned cell embeddings integrate the optimization of ℒ_CO_ and ℒ_PC_, utilizing the gene distribution at subcellular resolution alongside the cellular relative distribution at the cellular resolution. This joint approach allows for the extraction of cell embeddings from various levels of information, capitalizing on complementary insights. Subsequently, these heterogeneous losses are aggregated to form the total loss function, denoted as ℒ_total_.

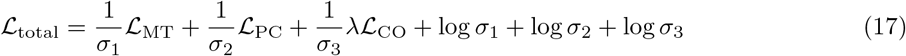

To avoid the domination of a certain task, we use the dynamic weight technique, which weights different losses with the homoscedastic uncertainty [37], enabling the weights to change dynamically during the training. {σ_1_, σ_2_, σ_3_} denotes the uncertainty weights for different tasks. λ is used as a linear decrease to force the model to focus less on the easier supervised learning tasks, λ is 1 from the beginning of the training and decreases to nearly 0 at the end of training. We believe that this training scheme can be treated as an inductive knowledge transfer which improves the generalisation of the model by sharing the domain information between complementary tasks.

### 4.9 SpatialFormer pretraining

SpatialFormer is a large transformer encoder-based model that utilizes a multi-task learning strategy to enhance the spatial awareness of cell embeddings through intertwined gene-gene co-occurrence and cell-cell co-localization within the tissue context. Our multi-task pre-training scheme captures hierarchical information about cells by focusing on both gene-gene interactions and cell-cell spatial relationships. This approach allows the model to gather multifaceted information across various levels, making it potentially applicable to a diverse range of downstream tasks.

In the masked tokens prediction, the model needs to figure out the expression order of genes by leveraging the surrounding context. The masked strategy is inherent in BERT, which sets 15% of tokens to be masked using the `<MASK>` token during training as 𝒰_mask_. To improve robustness, similar to the approach used in BERT [11], the masked tokens are treated as follows: 80% are actually masked, 10% are replaced with random tokens, and the remaining 10% are left unchanged. Special tokens, including `<CLS>`, `<SEP>`, `<PAD>`, are skipped in the masking stage.

To enable the multi-tasks pretraining, which updates the model parameters for different aspects, one single batch is used for training all the MT, PC, and CO tasks. An additional token classification head and pair head are applied to the final embeddings to predict the original identities of the masked tokens and cell pairs. Gene co-occurrence prediction is achieved by an adjacency projector layer, which outer products the 1D cell embeddings to 2D with the gene adjacency pair information. All the tasks are trained consecutively with the balanced weight scheme (**see 4.8**).

SpatialFormer was pre-trained with SpatialPC-300M dataset on the LUMI cluster using 64 AMD MI250 GPUs in 8 nodes with automatic mix precision as bfloat16 for 14 days. Each GPU loads the different cell pairs from different slides to enable randomness. AdamW Optimizer was used, with both *β*_1_ and *β*_2_ set to 0.9 and a weight decay of 0.1. The batch size was 1024, and the learning rate was initially set at 1 ×10^∈5^ and increased to 1 ×10^∈3^ over a warm-up period of 10,000 steps. Gradient clipping was applied at 1 to prevent gradient explosion. The model was trained for 100,000 steps until convergence, ensuring that the loss was minimized across all iterations

### 4.10 Gene pair differential analysis

Analogous to the discovery of differentially expressed genes, gene pairs that exhibit spatial co-occurrence can be identified by calculating the log fold change using multiple hypothesis testing. Genes assigned to a co-localized label can be encoded as gene pairs within specific cells. Cells containing at least one gene pair are filtered to construct a gene co-occurrence matrix, and the top 2000 highly variable gene pairs are selected for further analysis. Subsequent expression analyses can be performed based on predefined cell types. We utilize Scanpy [38] to load the data and conduct statistical tests, facilitating the identification of cell-type-specific differentially expressed gene pairs for downstream analysis.

### 4.11 Evaluating model Zero-shot performance

#### 4.11.1 Batch correction

To test whether SpatialFormer has learned robust features from the expression data, we employed the pre-trained model to perform inference on an unseen single-cell dataset, assessing the zero-shot capability of our model. We conducted the analysis using the integrated COVID-19 dataset [24], which consists of 18 batches of single-cell data.

The embeddings can be generated using the developed function `sp.tl.embed data` (https://github.com/TerminatorJ/Spatialformer/tree/master/spatialformer/tools), which integrates seamlessly with the AnnData object when using scanpy, the most popular single-cell analysis toolkit in python community. This function supports both single-cell and pair-wise cell inputs, automatically tokenizing the genes and constructing the collator to facilitate model execution.

#### 4.11.2 Cell-cell interaction prediction

Testing whether any two cells are co-localized is a fundamental task in revealing the cell composition and cell density of the cellular niches. The sparse binary adjacency matrix we built previously (**see 4.5**) can directly infer the constitution of cellular niches. The query cell *i* can be treated as the centroid cell in the niches with radius r. key cells within the distance of r are the connected neighbors. Therefore, we can get the niche composition of cell type *k* as 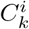

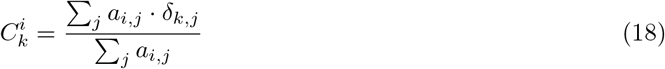

where *a*_*i*,*j*_ = 1 indicates a connection between cell *i* and cell *j*, while *δ* is the indicator function which indicates whether the cell type of cell *j* belongs to cell type *k*. According to the cell type distribution of each query cell, we can calculate the distribution of all cell types in a slide. However, it is computationally unrealistic to perform inference across all cellular combinations to obtain binary predictions, so we randomly selected 500 cells for evaluation. The divergence of the real cell type distribution can be compared with the predicted cell type distribution 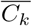.The Jensen-Shannon divergence can be used to test the differences between two distributions.

The Jensen-Shannon divergence (JSD) between two probability distributions *P* and *Q* is defined as follows:

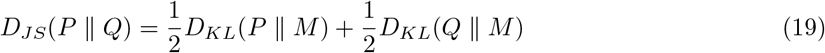

where 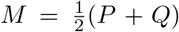 is the average of the two distributions, and *D*_*KL*_ is the Kullback-Leibler divergence, defined as:

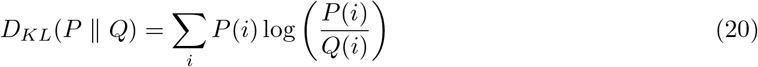

### 4.12 Fine-tuning the model

#### 4.12.1 Cell identities identification

Instead of being trained with paired cells, SpatialFormer is fine-tuned to adapt to single input sequences, enabling it to handle tasks such as cell type annotation. The SpatialCC-12M dataset is utilized to adapt the model to a single input mode. The training objective incorporates only the masked token prediction (MT) and co-occurrence (CO) tasks, formulated as:

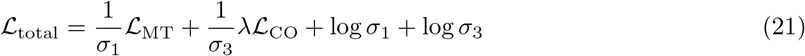

The model is trained for an additional 100,000 steps to enhance its understanding of single input data. Subsequently, the [CLS] token is retained to derive cell-level embeddings.

Following this, the model was frozen, and a simple multi-layer perceptron (MLP) was employed as a probe network to predict cell type labels. Early stopping was implemented to prevent overfitting. The predicted labels were then projected back onto the original PCA UMAP for improved visualization of prediction accuracy.

Further pre-training at the slide level employs three specific optimization objectives as formulated (18). The model underwent an additional 50,000 steps of pre-training to enhance slide-specific learning.

### 4.13 Perturbation analysis

To analyze the importance of specific gene pairs in gene-gene co-localization, we implemented Iterative Gene Pair Perturbation (IGP) to systematically knock out gene pairs from the pair-wise combinations of ranked gene tokens. Let *L*_1_ represent the number of genes in cell A and *L*_2_ the number of genes in cell B. We manually deleted each gene pair in a total of *L*_1_ × *L*_2_ iterations while retaining the other genes to generate new predicted logits. The Prediction Class (PC) head was employed to compute the deduction *d* between the original logits and perturbed logits. To assess the presence of interactions across slides for any two cell types of interest, we conducted a one-sample one-tailed t-test based on the standard Gaussian distribution across all cells belonging to these two cell types. Namely, we are testing if the observed *d* for a certain cell type pair has a mean value significantly lower than 0. This test evaluated the null hypothesis that the parameter *d* does not significantly affect the interaction between the two cell types in relation to the perturbed gene pairs.

---

**Supplementary Figure 1.**
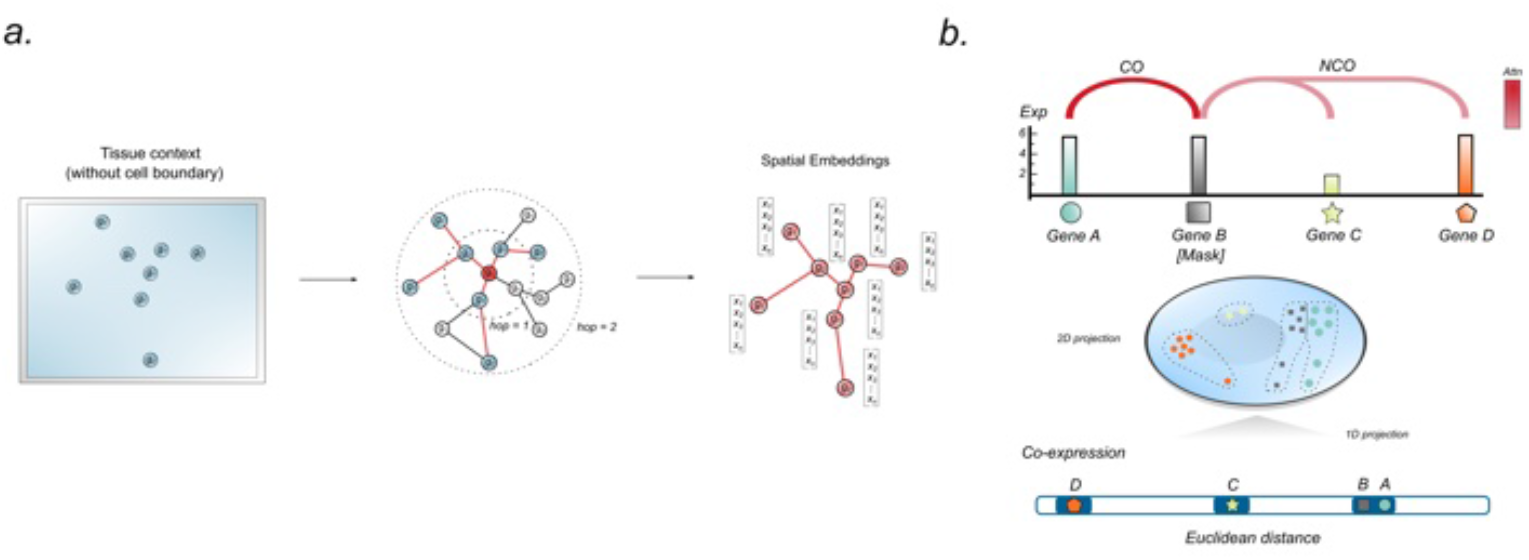
Characterization of abundance spatial gene features. **a**, Global gene-gene interactions generated by GraphSAGE. Transcript embeddings are derived by constructing connected neighbors from randomly selected transcripts within a two-hop distance. These embeddings can be utilized to systematically investigate the overall scale of these relationships, thereby enhancing the model’s pre-training process. **b**, Characteristics of Regional Gene Spatial Features: Rather than exclusively focusing on expression levels, genes with similar expression levels can be further classified based on their spatial distances within cells.

**Supplementary Figure 2.**
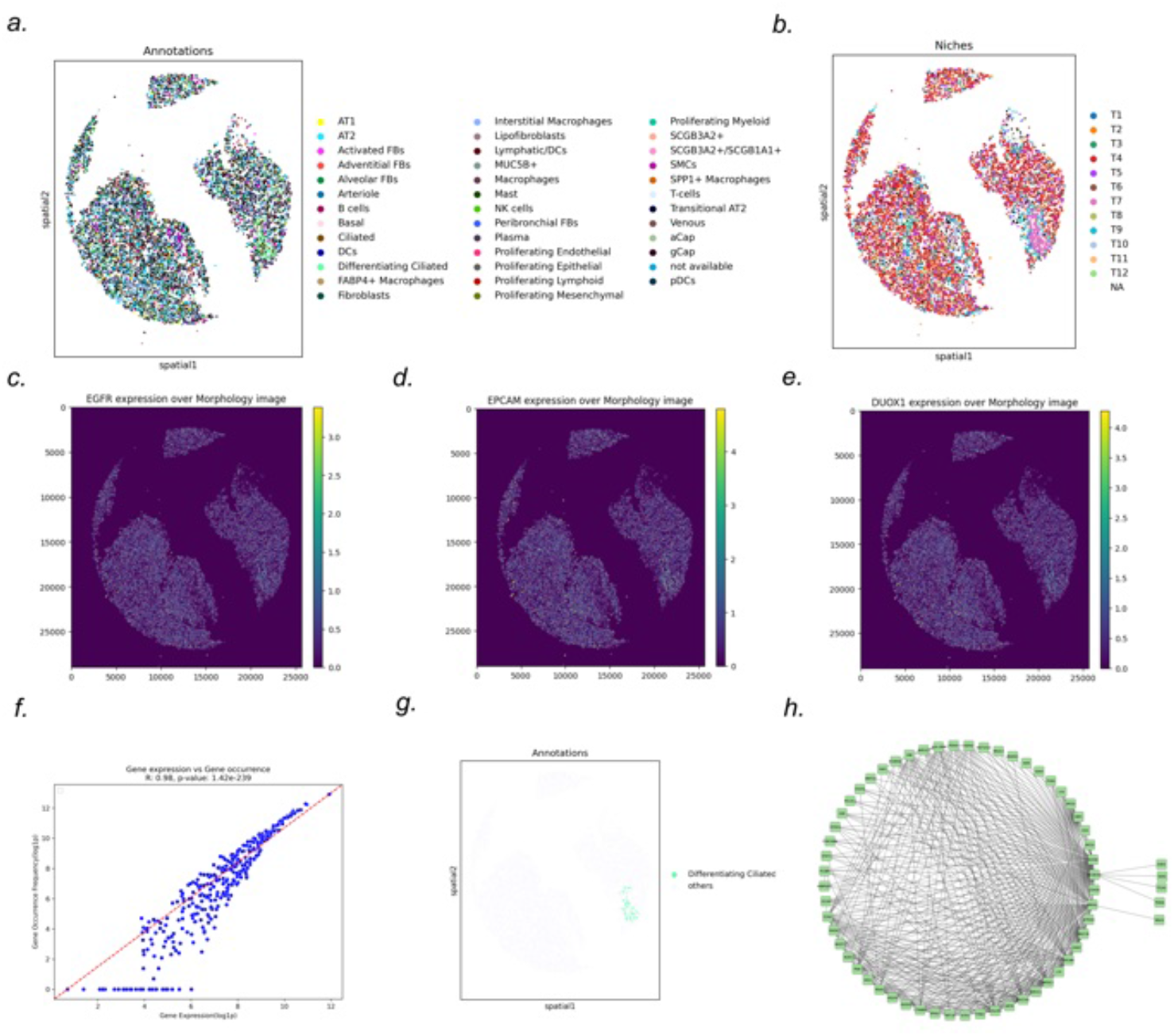
Gene co-occurrence across the tissue. **a-b**, Cell type and niche annotations with the spatial context in slide THD0008. **c-e**, Visualization of the expression levels of EGFR, EPCAM, and DUOXI overlaid on the morphological image. **f**, Correlation between gene expression levels and their frequency in slide THD0008. **g**, Annotation of differentiating ciliated cells within the spatial context. **h**. Gene-gene interaction network for slide THD0008, visualized using Cytoscape.

**Supplementary Figure 3.**
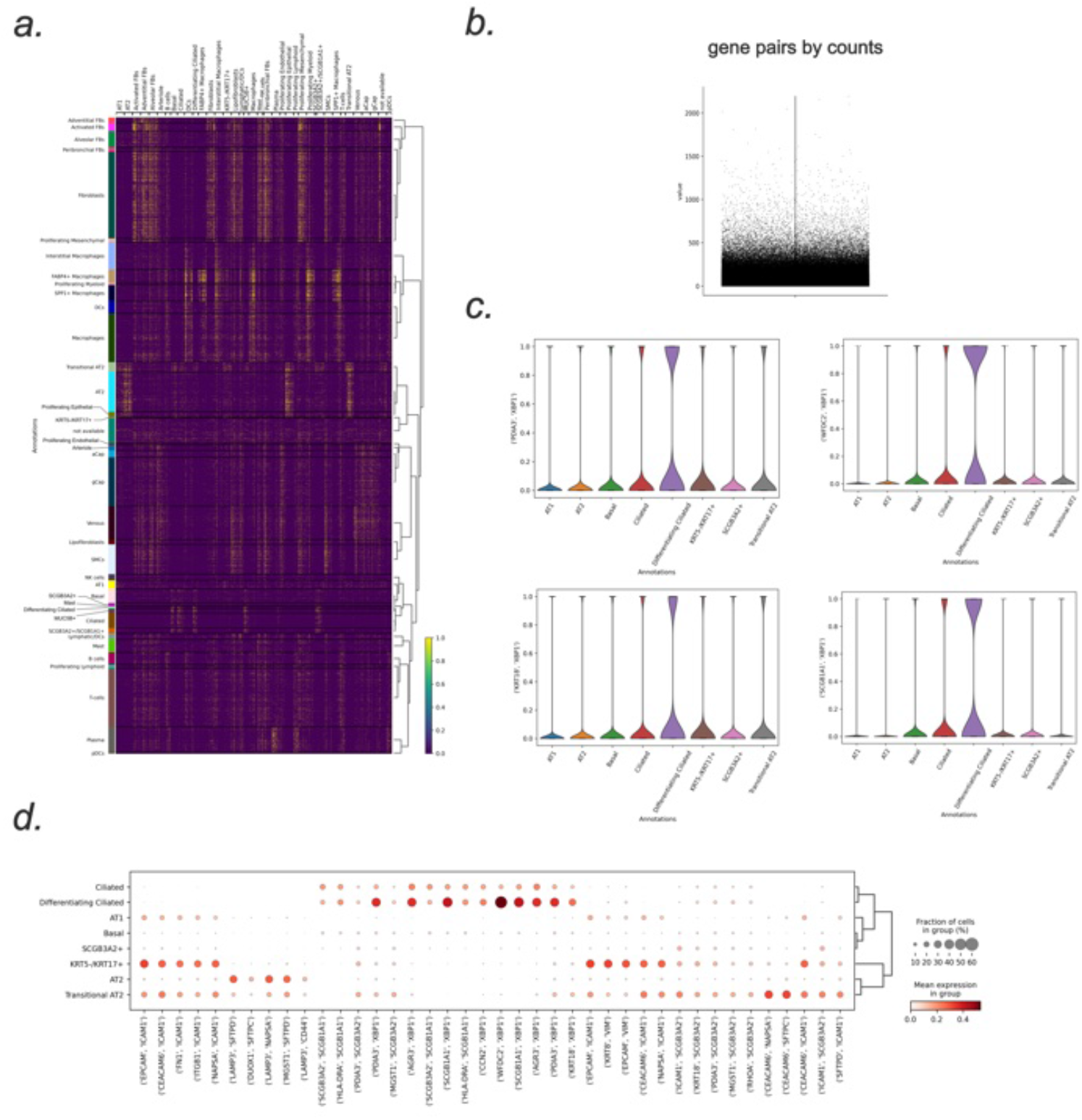
The differential expressed gene pair across cell types. **a**. Heatmap showing the binary gene pair markers across all the cell types. Rows correspond to specific cell types, while columns reflect co-localized pairs appear in cell type. **b**. violin plot illustrating the distribution of gene pairs by counts in pulmonary lung dataset. **c**. Distribution of four top different gene pairs for differentiating ciliated. **d**. Dot plot depicting the selected significant gene-gene co-occurrence pairs for 8 different cell types.

**Supplementary Figure 4.**
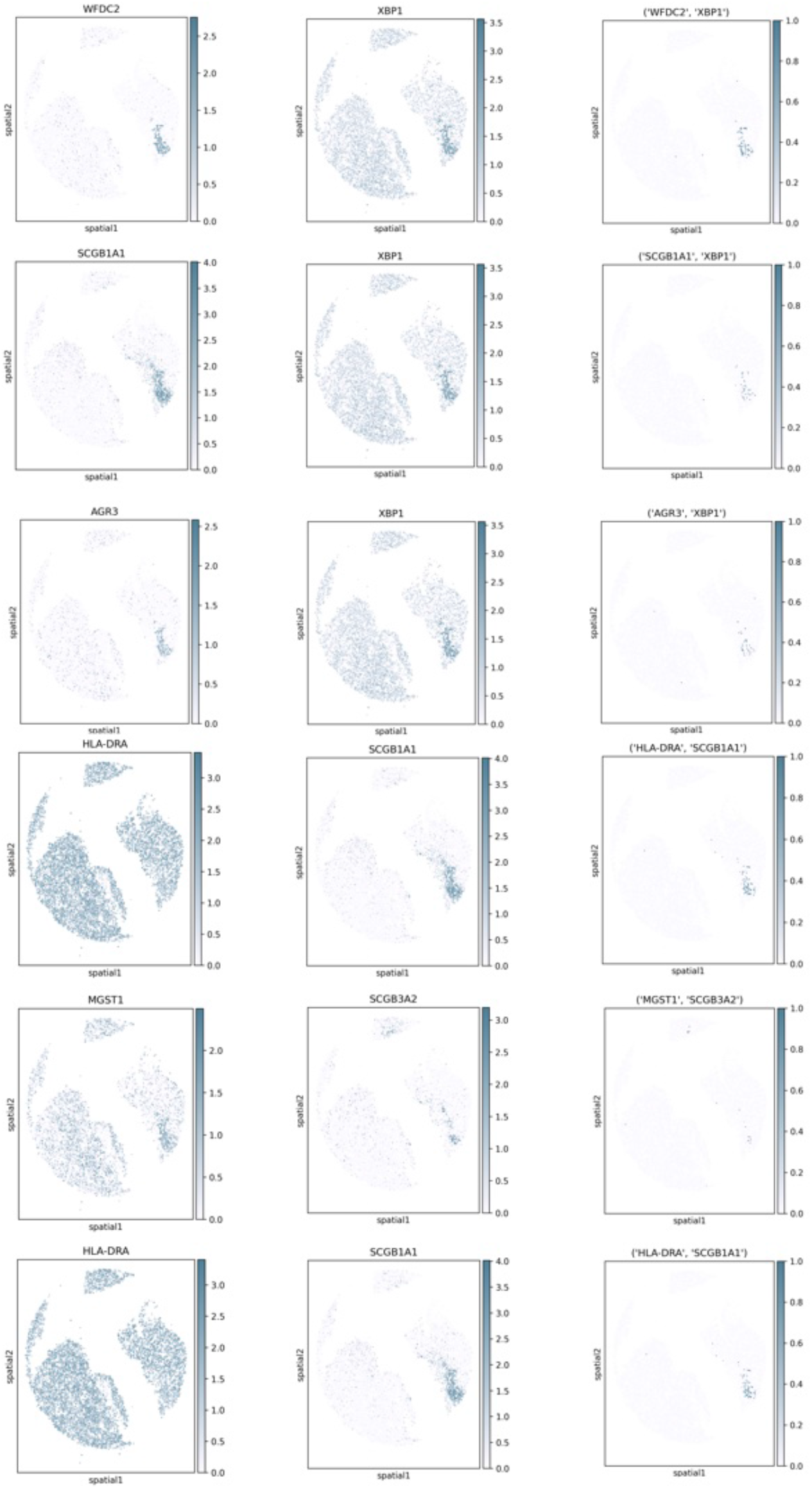
The co-occurrence of gene pairs enhances the sensitivity of cell type identification. The two left columns display the expression levels of individual genes within the spatial context. The far-right column indicates the binary expression of spatially co-occurring gene pairs, with cells colored only if the gene pairs co-localized within a single cell.

**Supplementary Figure 5.**
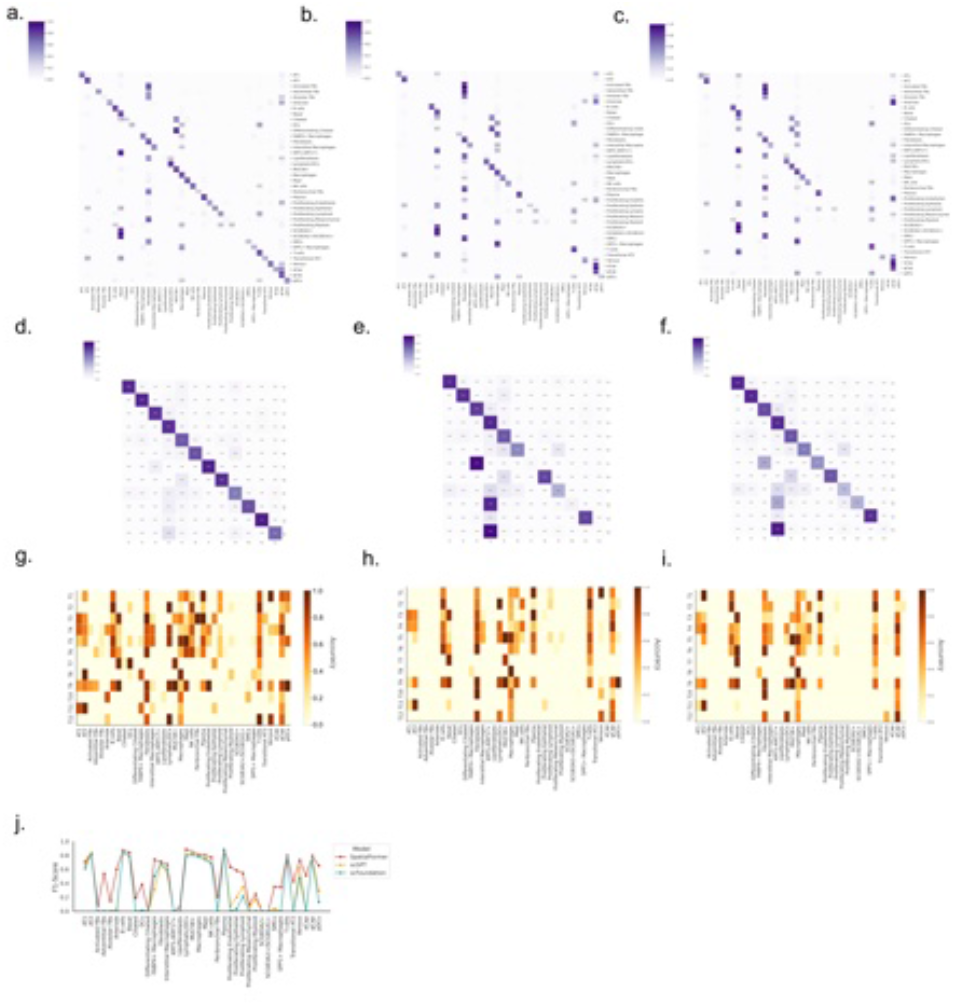
Prediction results of the probe network across different cell types and niches. **a-c**, Heatmaps illustrating the true positive of different cell types across various language models. The panels, from left to right, represent SpatialFormer, scGPT, and scFoundation, respectively. **d-f**, Heatmaps showing the accuracy among different cell niches for the same language models, with panels organized similarly. **g**, A line plot depicting the F1 score across different cell types, with distinct colors representing different language models. **h-i**, Heatmaps indicating accuracy scores across cell niches and types, where darker cells signify higher accuracy scores. All evaluations are based on 10% of cells from the VUILD110 sample.

**Supplementary Figure 6.**
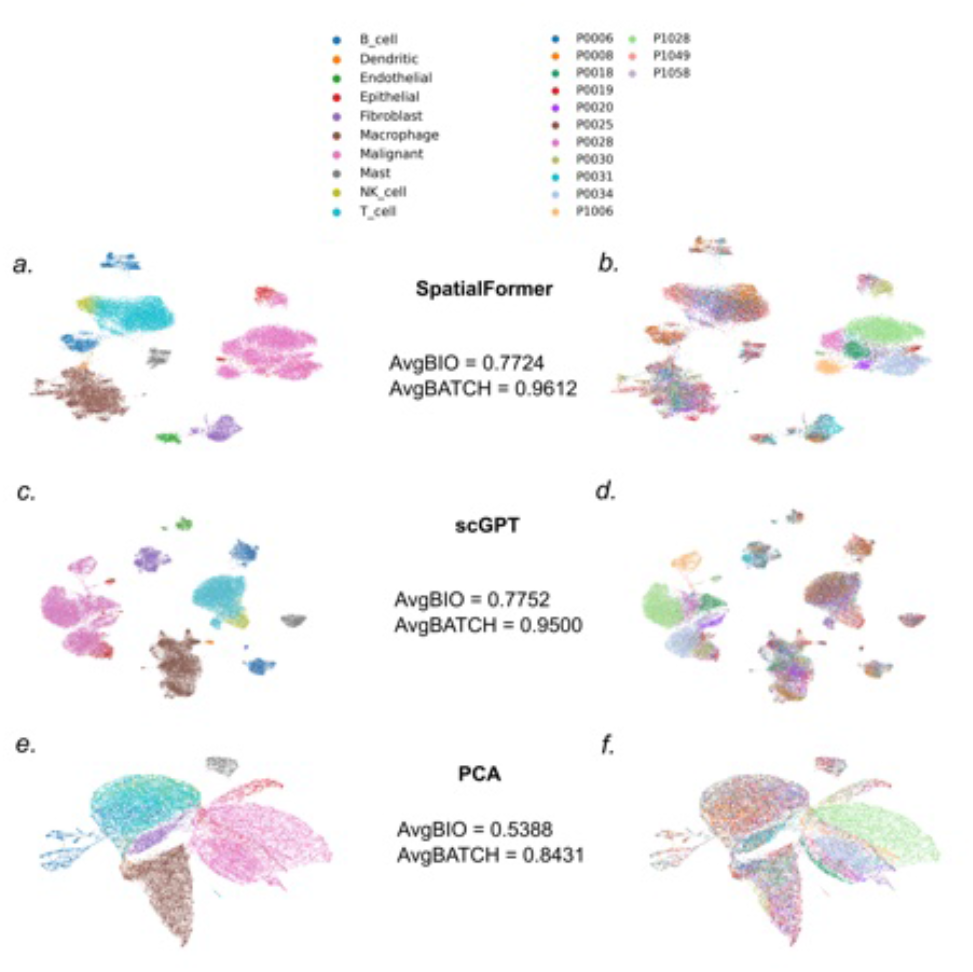
Performance of Batch Integration for the Kim Lung Dataset. **a-b**, Embedding results from SpatialFormer, with cells annotated by cell types in the left panel and by batch in the right panel. **c-d**, Batch integration results from scGPT, similarly labeled with cell types on the left and batch on the right. **e-f**, Batch integration results from PCA, with cell annotations indicating types on the left panel and batch information on the right. The color bar is aligned at the top, corresponding to both the left and right panels of the plot.

**Supplementary Figure 7.**
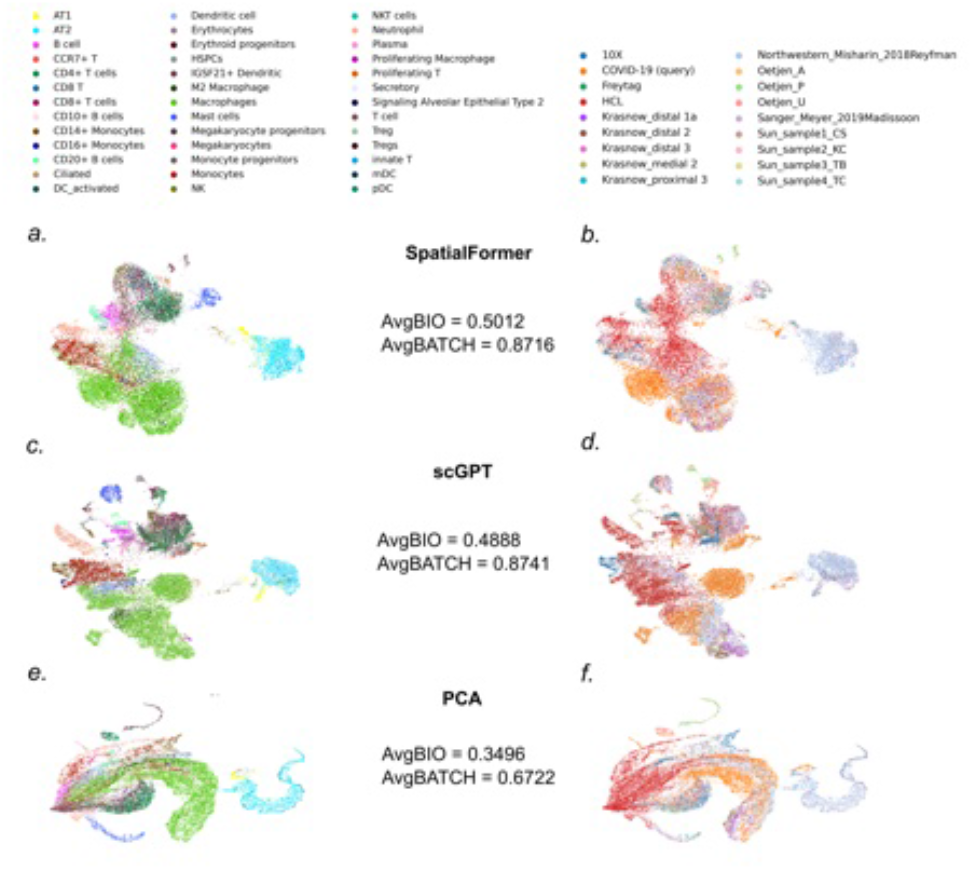
Performance of Batch Integration for the COVID-19 Dataset. **a-b**, Embedding results from SpatialFormer, with cells annotated by cell types in the left panel and by batch in the right panel. **c-d**, Batch integration results from scGPT, similarly labeled with cell types on the left and batch on the right. **e-f**, Batch integration results from PCA, with cell annotations indicating types on the left panel and batch information on the right. The color bar is aligned at the top, corresponding to both the left and right panels of the plot.

**Supplementary Figure 8.**
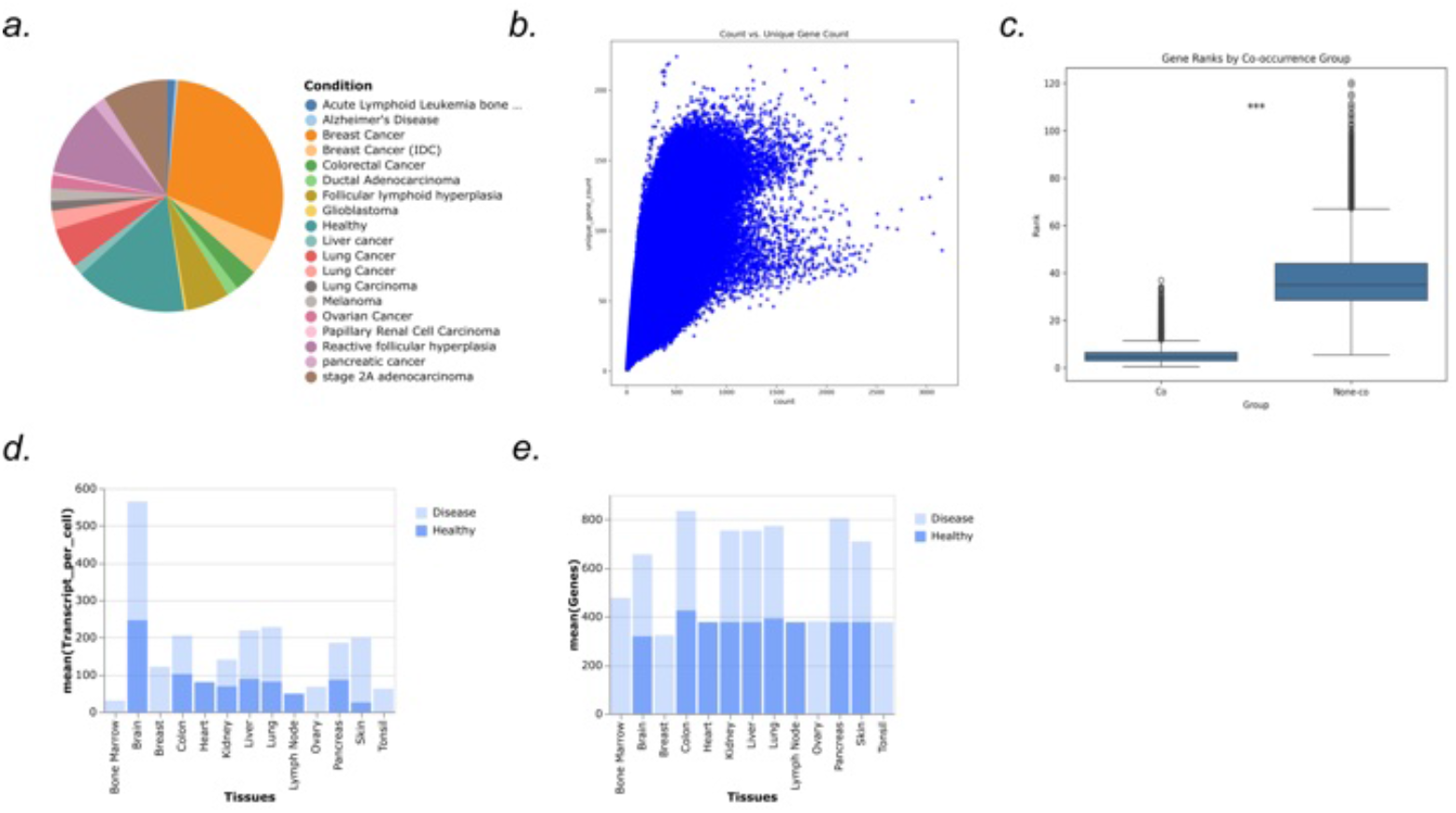
The overview of the pre-training dataset. **a** Proportions of healthy and diverse disease samples. **b** Statistical distribution of sequence lengths, where the y-axis represents the input sequence length and the x-axis denotes the cumulative count of lengths. **c** Rank distribution of co-occurring versus non-co-occurring genes. **d-e** Mean number of transcripts and genes across different tissues within disease and healthy categories.

**Supplementary Figure 9.**
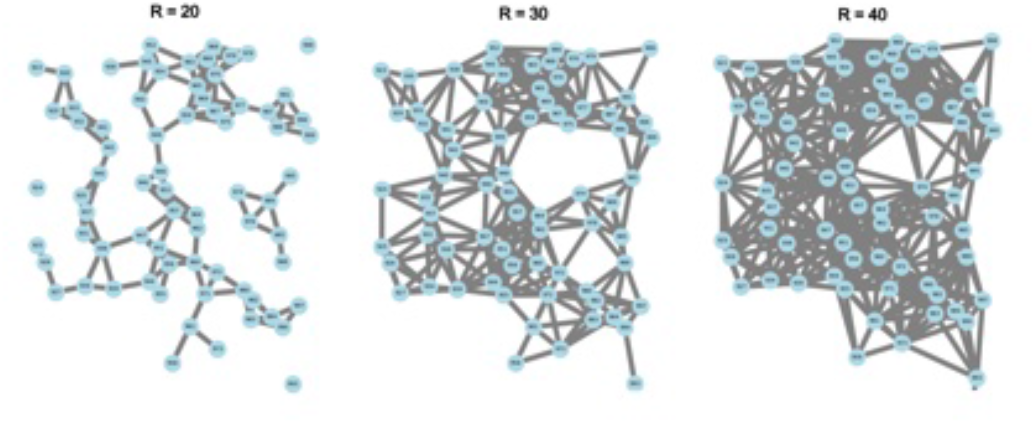
The cell connection graph of different radius options.

